# Static magnetic stimulation induces structural plasticity at the axon initial segment of inhibitory cortical neurons

**DOI:** 10.1101/2022.01.26.477963

**Authors:** J.L. Beros, E.S. King, D. Clarke, J. Rodger, A.D Tang

**Affiliations:** The University of Western Australia, School of Biological Sciences, Crawley, Western Australia; The Perron Institute for Neurological and Translational Science, Nedlands, Western Australia; The University of Montreal, Montreal, Quebec; The University of Western Australia, School of Biomedical Sciences, Crawley, Western Australia

## Abstract

Static magnetic stimulation (SMS) is a form of non-invasive brain stimulation that can alter neural activity and induce neural plasticity that outlasts the period of stimulation. While SMS is typically delivered for short periods (e.g., 10 minutes) to alter corticospinal excitability or motor behaviours, the plasticity mechanisms that can be induced with longer periods of stimulation have not been explored. In mammalian neurons, the axon initial segment (AIS) is the site of action potential initiation and undergoes structural plasticity as a homeostatic mechanism to counteract chronic changes in neuronal activity. Therefore, we investigated whether the chronic application of SMS would induce structural AIS plasticity in cortical neurons. SMS (0.5 Tesla in intensity) was delivered to postnatally derived mouse primary cortical neurons consisting of mainly inhibitory neurons, for 6 or 48 hours beginning from 7 days *in vitro* (DIV7). AIS structural plasticity (length and starting distance from the soma) was quantified immediately after and 24 hours post-stimulation. Following 6 hours of stimulation, we observed an immediate decrease in median AIS length compared to control, that persisted to 24 hours post stimulation. In addition, there was a distal shift in the AIS start position relative to the soma that was only observed 24 hours after the 6-hour stimulation. Following 48 hours of stimulation, we observed an immediate shortening of AIS length and a distal shift in AIS start position relative to the soma, however only the distal shift in AIS start position persisted to 24 hours post-stimulation. Our findings provide the foundation to expand the use of SMS to more chronic applications as a method to study or promote AIS plasticity non-invasively.

## Introduction

Non-invasive forms of brain stimulation that modulate neural activity or excitability have great potential as a tool to study neural plasticity mechanisms and as a treatment for neurological disorders. In particular, static magnetic stimulation (SMS) is a form of non-invasive brain stimulation where static magnetic fields are delivered to neurons by a rare-earth neodymium magnet placed at the surface of the skull (1). In studies using human participants, SMS has been shown to modulate neural activity in the motor and sensory cortices (2–7). In particular, delivering SMS for short periods (e.g., 10-30 minutes), most commonly leads to a reduction in cortical excitability and activity (3, 4, 6–9), which may suggest that SMS leads to an acute increase in neural inhibition (5, 8). While many studies have focused on SMS and its immediate effects, more chronic applications of this intervention and its effect on neural plasticity and excitability is not known.

The axon initial segment (AIS) is the microdomain of the axon containing an abundance of ion channels and anchoring and structural proteins that facilitate action potential initiation (10). It is well established that the AIS of individual neurons undergoes structural remodelling (i.e. structural plasticity) in response to significant periods of high and low neural activity. Specifically, AIS structural plasticity includes changes in AIS length and starting position along the axon (10, 11). These changes in AIS structure are believed to be a homeostatic response to perturbations in neural activity, as they have a direct functional consequence on neuronal excitability (10–12). For example, *in vitro* and *in vivo* models have shown that significantly increasing neural activity via pharmacological or optogenetic interventions, leads to a shortening of the AIS and/or a relocation to more distal start positions along the axon and vice versa for significant decreases in neural activity (12–14).

Given that SMS is a simple method that can be used to alter neural activity, we sought to determine whether chronic SMS could be used to induce structural AIS plasticity of cortical neurons. We characterised the structure of the AIS of cortical neurons *in vitro*, following 6 or 48 hours of SMS. Mouse primary cortical neurons dissected from postnatal day 1 (P1) were exposed to SMS (0.5T) and processed for immunofluorescence to determine AIS length and starting position along the axon. We found that SMS induced an AIS shortening and/or distal relocation in inhibitory neurons after 6 and 48 hours of stimulation, with some of these changes persisting to 24 hours post-stimulation. Interestingly, the extent of AIS structural plasticity was similar to the AIS structural plasticity induced with the chronic application of 15mM potassium chloride (KCl), which is commonly used to induce structural plasticity *in vitro*. Overall, our results suggest that chronic SMS induces structural plasticity in inhibitory cortical neurons and could be a promising tool to induce AIS plasticity non-invasively.

## Methods

### Preparation and maintenance of cortical cell cultures

P1 C57Bl/6J mice were delivered from the Animal Resource Centre (Murdoch, Western Australia) to the University of Western Australia. Animals were euthanised with an overdose of sodium pentobarbitone (Lethabarb, Virbac) administered by an intraperitoneal injection. Animal procedures were approved by the UWA Animal Ethics Committee prior to commencement (RA/3/100/1677). After confirmation of death (absence of pinch reflex), the cortices of each mouse were dissected into medium consisting of 2% B27 plus supplement (Gibco), 0.25% GlutaMAX supplement (Gibco) in Hibernate A medium (Gibco). The meninges were carefully removed from the dissected cortices, which were then cut into smaller pieces (approx. 0.5mm^2^) using a scalpel blade. Cortices were equilibrated at 30°C for 10 minutes prior to transferring them into pre-warmed dissociation medium consisting of 2mg/ml papain (Sigma-Aldrich) and 0.25% Glutamax (Gibco) in Hibernate A medium without calcium and magnesium (BrainBits) that was pre-filtered through a 0.2μm filter. This mixture was spun at 350 rpm at 30°C for 30mins, after which, the cortices were transferred into 2ml of pre-warmed dissection medium. Cortices were triturated using a sterile glass Pasteur pipette (Corning) and the supernatant was transferred into a 15mL centrifuge tube. After repeating the trituration process and supernatant removal for an additional two times, the collected supernatant was centrifuged at 12,000 rpm for 5 minutes. The supernatant was aspirated and the pellet was resuspended in culture medium consisting of 2% B27^+^ supplement (Gibco), 0.25% Glutamax (Gibco) and 0.1% penicillin/streptomycin (Gibco) in Neurobasal Plus medium (Gibco). Cell counts were estimated using a haemocytometer and seeded onto 13mm diameter coverslips in 24 well plates (Trajan) pre-coated with 50μg/ml poly-d-lysine (Gibco) and 40μg/ml mouse type 1 laminin (Gibco) at a seeding density of 150,000 cells/coverslip. Cell culture plates were incubated at 37°C with 5% CO_2_ for a period of 7 days before stimulation, with 50% of the media replaced at 4 days *in vitro* (DIV4).

### Stimulation of cell cultures

On the morning of DIV7, 50% of the cell culture medium was replaced from each well prior to treatment with pharmacological (positive control), static magnetic or control stimulation (negative control). Pharmacological stimulation was achieved by dissolving 15mM of potassium chloride (KCl) into the cell culture medium, previously shown to increase neuronal activity in culture and induce morphological changes in the AIS within 6 hours of stimulation (13). SMS was delivered by mounting the culture plate and coverslip on a north-facing 13mm diameter rare earth neodymium magnet (0.5T intensity at surface of the magnet) within the incubator. Control cultures did not receive any stimulation protocol and were maintained in normal culture medium in the incubator for the stimulation times. To assess the immediate effects of our stimulation protocols, the stimulation interventions were applied for 6 or 48 hours and fixed immediately post-stimulation. To assess any delayed effects following stimulation, after completion of the 6 and 48 hour stimulation periods, 90% of the cell culture medium was replaced in each well and cells were fixed 24 hours later. Prior to fixation, cell culture medium was removed from all wells, washed twice with sterile phosphate buffered saline (PBS; Gibco) and fixed with 4% paraformaldehyde in 0.1M PBS for 8 minutes. After the fixation period, wells were washed twice with 0.1M PBS and stored in 0.1M PBS with 0.1% sodium azide.

### Immunohistochemistry

Cell culture wells were washed twice with 0.1M PBS followed by permeabilsation with 0.2% Triton-X in 0.1M PBS for 10 mins. Following this, cultures underwent a blocking step with 10% normal goat serum (ThermoFisher) in 0.2% Triton-X in PBS for 1 hour. To identify neurons, cells were immunolabelled for microtubule-associated protein 2 (MAP2), expressed in the soma and dendrites of neurons, in addition to glutamic acid decarboxylase 65/67 (GAD65/67), a marker of inhibitory neurons expressed in the soma and processes. For AIS analysis, we paired the neuronal markers with antibodies against ankyrin G (AnkG), an anchoring protein highly expressed at the AIS and commonly utilised to identify the AIS and AIS structural plasticity (12–15). Cultures were incubated in primary antibodies for mouse anti-AnkG (1:500, Thermofisher; catalogue number 33-8800), rabbit anti-GAD 65/67 (1:1000, Thermofisher; catalogue number PA5-36080) and chicken anti-MAP2 (1:5000, Abcam; catalogue number ab5392) in a solution of 2% normal goat serum in 0.2% Triton-X in 0.1M PBS at 4°C for 24 hours. After primary antibody incubation, wells were washed three times with 0.1M PBS and then incubated with secondary antibodies goat anti-mouse Alexa Fluor 488 (1:600, Invitrogen), goat anti-rabbit Alexa Fluor 594 (1:600, Invitrogen) and goat anti-chicken Alexa Fluor 647 (1:600. Invitrogen) in the same primary blocking solution for 2 hours at room temperature. The nuclear label Hoechst (1:1000, Invitrogen) was included in the secondary antibody incubation as a non-specific cell marker. Following this, wells were washed three times with 0.1M PBS and coverslips were mounted on glass slides (Hurst Scientific) with Prolong^™^ diamond antifade mountant (Life Technologies).

### Imaging and cell quantification

After a minimum of 48 hours, coverslips were imaged on a C2 confocal microscope (Nikon) at 60x magnification (oil immersion, NA=1.40, Nikon Plan Apo). Images were acquired in NIS Elements (Nikon) at a resolution of 512 x 512 pixels. The total area of the coverslip was divided into thirds and 15 fields of view were acquired in a systematic random manner (5 fields of view from each third). This represented a total sample area of approximately 0.5% of each coverslip. Multiple images were captured in the z-plane of each field of view, at a z-step depth of 0.125um, capturing the appearance and disappearance of all AIS in the field of view. Images were processed in FIJI (Image J) to generate a single RGB TIFF of the maximum z projection. Prior to analysis, investigators were blinded to the stimulation group that the images belonged to.

AIS length was quantified using a custom Matlab code developed by Grubb and colleagues (13). Briefly, the axon is manually traced using the AnkG fluorescence intensity, with the start and end position of the AIS defined as the first and last point along the axon where the AnkG fluorescence profile diminished to 0.33 of the maximum fluorescence value (16). Cells that showed branching of the AIS/AnkG were excluded from the analysis. The start position of the AIS relative to the soma was estimated using Fiji image processing software by measuring the distance between the end of the MAP2 fluorescence signal and the start of AnkG fluorescence. Cells were classified as inhibitory neurons if they showed positive co-labelling for MAP2, GAD65/67 and DAPI.

### Statistical analysis

The data collected from each neuron was considered an independent sample, however we also report the number of coverslips and cultures that the data consists of. Both AIS length and distance from the soma were analysed with Kruskall-Wallis tests with follow up Dunn’s post-hoc comparisons. The data is reported as median ± SEM. Results were considered statistically significant at α <0.05 and additional cumulative frequency distributions were plotted to aid in the interpretation of our results (see supplementary Figure 1).

## Results

Similar to previous studies, the AIS was identified by the immunofluorescence profile of AnkG (see Figure 1). We observed heterogeneity in AIS morphology in our cell cultures (17), with the AIS originating both immediately proximal and more distal to the soma, indicating that our cultures contained a mixture of cells that had dendritic and somatic originating axons. Surprisingly, the majority of neurons present in our cell cultures at our assessed ages were immunopositive for the inhibitory marker GAD65/67 (mean of 83%, quantified from 5 control media coverslips from 5 culture experiments). In the data we collected for AIS analysis, only a small population of MAP2 positive neurons did not express the inhibitory marker (134/1544 neurons) and therefore, we restricted our statistical analysis to GAD65/67 immunopositive neurons only.

**Figure 1.**
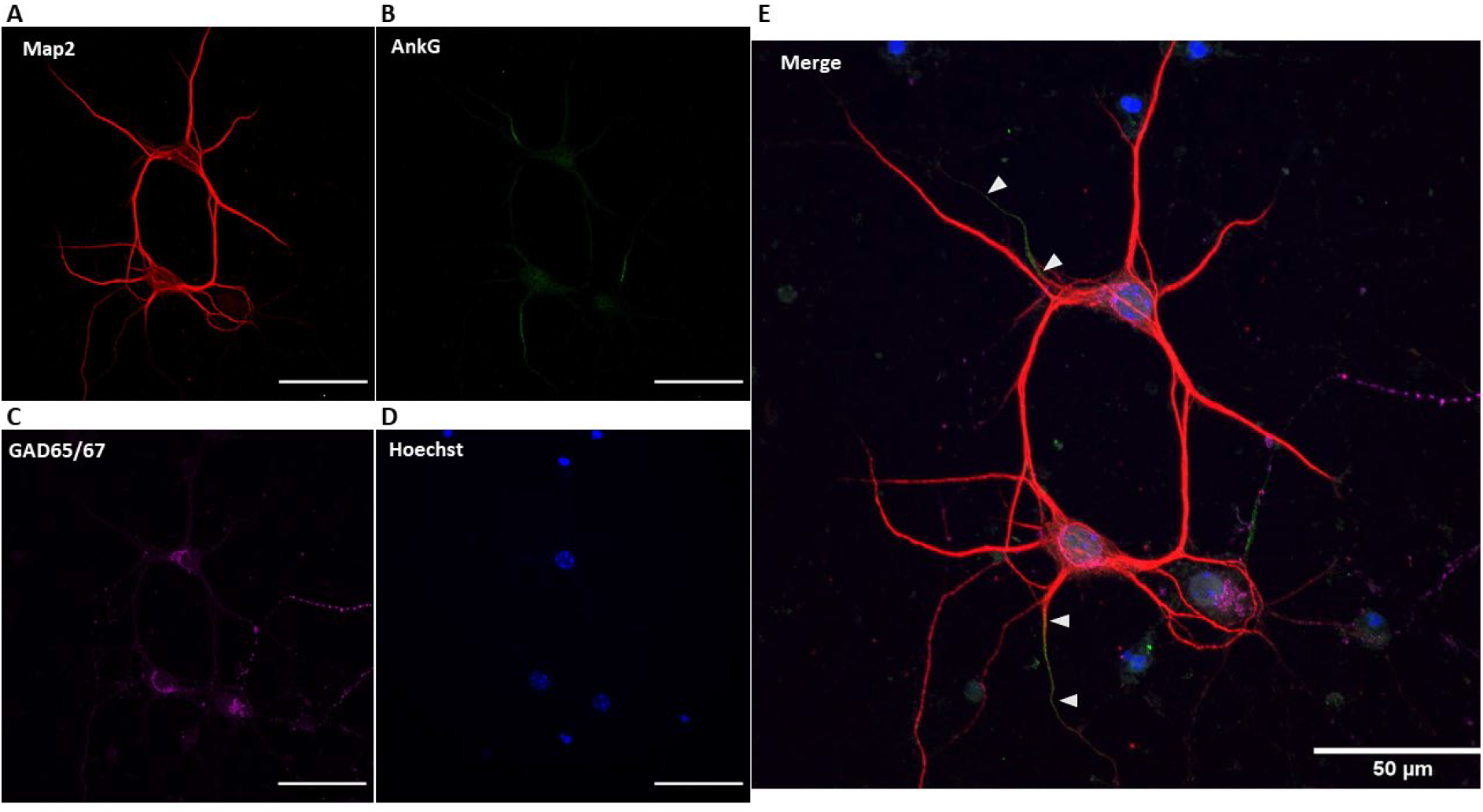
Representative maximum intensity z-projection of immunolabelled inhibitory cortical neurons in cell culture at DIV7. Neurons were immunolabeled for (A) the neuronal marker MAP2, (B) AIS marker AnkG, (C) inhibitory neuronal marker GAD65/67 and (D) nuclear marker Hoechst. Images were (E) merged and the AIS length and distance from soma was quantified. Triangles indicate the approximate start and end of the AIS. Images were captured on a C2 confocal microscope at 60x magnification. Scale bar = 50μm.

### KCl induced AIS structural plasticity in inhibitory neurons

Similar to previous AIS studies in embryonically derived primary neurons in culture, we found that the addition of 15mM KCl to the culture media induced a strong reduction in the length of the AIS in P1 derived cultures (see Figure 2). After incubation for 6 hours, the median length of the AIS was significantly reduced from 26.81±0.53μm in control conditions to 23.45±0.45μm with KCl (χ2(2) = 12.02, p < 0.01, see Figure 2A). A significant reduction in median AIS length was also present after 48 hours of stimulation (χ2(2) = 12.72, p <0.01), reducing from 26.08±0.44μm in control conditions to 23.67±0.58μm with KCl (see Figure 2B). This represented a reduction in the length of the AIS of 13% at 6 hours and 10% at 48 hours. Interestingly, when cell cultures were returned to control media immediately after stimulation for 24 hours, the median AIS length in these conditions remained significantly shortened when compared to neurons in the control condition (27.05±0.51μm in control and 24.04±0.45μm in KCl; χ2(2) = 22.17, p < 0.001 at DIV8, Fig.2A; 23.31±0.42μm in control medium and 19.45±0.53μm in KCl stimulated neurons; χ2(2) = 22.21, p < 0.001 at DIV10, see Figure 2B).

**Figure 2.**
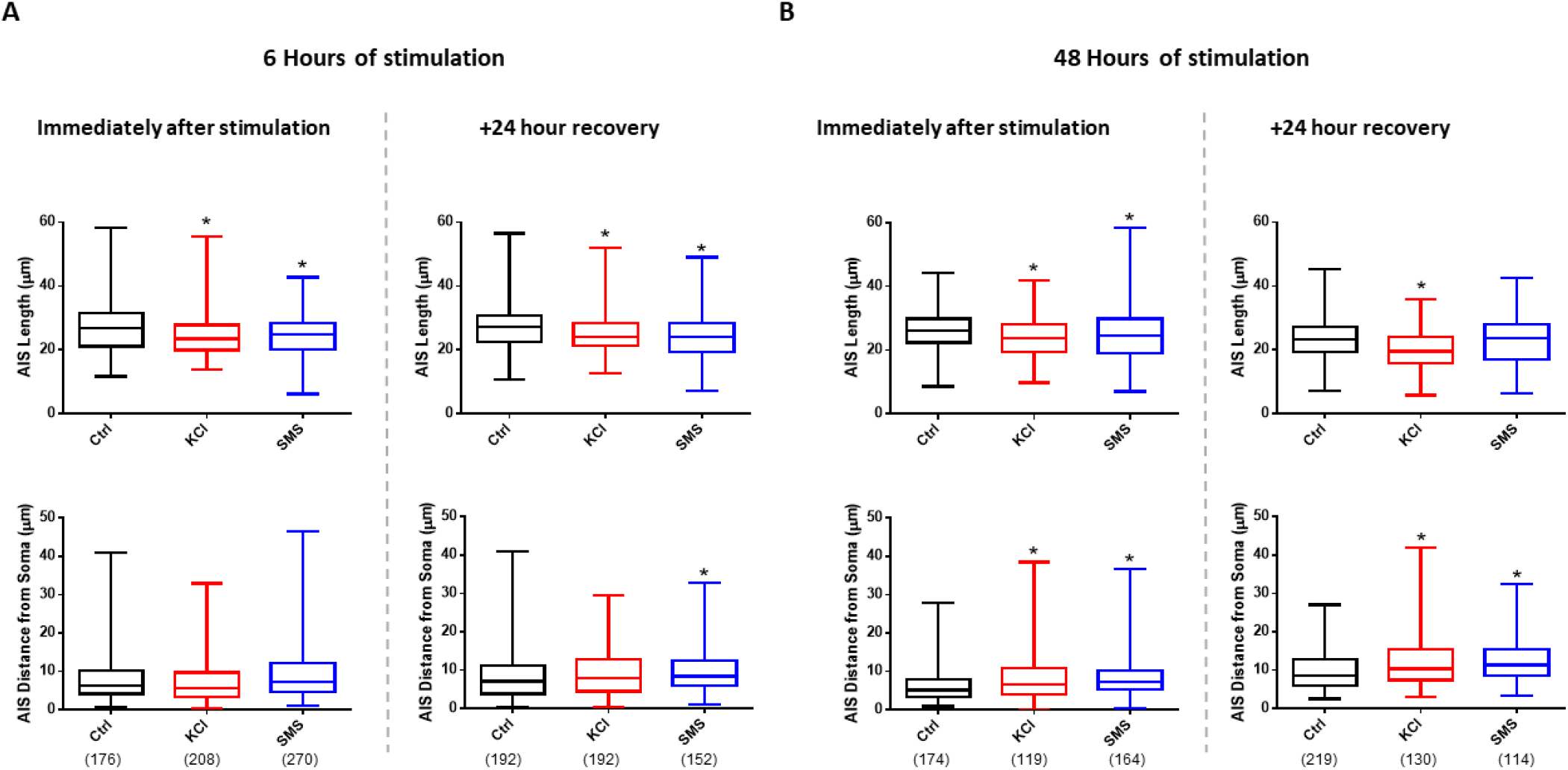
Quantification of AIS length and distance from soma. (A) Box plots of AIS length (top) and distance from soma (bottom) immediately after (left panel) and 24 hours after (right panel) 6 hours of stimulation. (A) Box plots of AIS length (top) and distance from soma (bottom) immediately after (left panel) and 24 hours after (right panel) 48 hours of stimulation. Number of neurons analysed in each group presented in brackets under the group names. Data from each treatment group at each DIV was collected from at least 5 coverslips from culture experiments, except the 48hr SMS +24 hours recovery group (2 cultures). * denotes p<0.05

In the cells that received 48 hours of KCl stimulation, the proximal start of the AIS was also located more distal when compared to control neurons (5.16±0.33μm in control medium and 6.64±0.68μm in KCl stimulated neurons; χ2(2) = 27.18, p < 0.001; see Figure 2B) and this significant difference persisted for 24 hours after removing the KCl and returning the neurons to control media (8.63±0.31μm in control medium and 10.31±0.64μm in KCl stimulated neurons; χ2(2) = 27.46, p < 0.001, see Figure 2B).

### SMS induced structural plasticity in inhibitory neurons

Similar to the 15mM KCl stimulation, neurons exposed to 0.5T SMS demonstrated a significant reduction in the median AIS length compared to neurons in the control condition (see Figure 2). Immediately after 6 hours of SMS, the median AIS length was significantly reduced (26.81±0.53μm in control medium and 24.86±0.41μm after SMS; χ2(2) = 12.02, p < 0.01, see Figure 2A) and this significant reduction was also observed immediately after 48 hours of SMS (26.08±0.44μm in control medium and 24.58±0.69μm after SMS; χ2(2) = 12.72, p <0.01, see Figure 2B). The length of the median AIS length remained significantly shorter 24 hours after the 6-hour stimulation had ended (27.05±0.51μm in control medium and 24.02±0.59μm after return from SMS; χ2(2) = 22.17, p <0.001, see Figure 2A). Additionally, when SMS was replaced with control medium for 24 hours following 48 hours of stimulation, the length of the AIS was not significantly different from control values (23.21±0.42μm in control medium and 23.53±0.69μm after SMS).

In addition to changes in the length of the AIS after SMS, there was a significant distal relocation in the start position of the AIS relative to the soma (see Figure 2). This was present immediately after 48 hours of SMS, increasing from a start location of 5.16±0.33μm from the soma in control medium to 7.19±0.41μm after SMS (χ2(2) = 27.18, p <0.001, see Figure 2B). Interestingly, there was a delayed distal relocation that was observed 24 hours after completion of SMS for 6 hours (7.01±0.46μm in control medium and 8.35±0.44μm after return from SMS; χ2(2) = 7.27, p <0.05, see Figure 2A) and 48 hours (8.63±0.31μm in control and 11.36±0.51μm in SMS; χ2(2) = 27.46, p <0.001, see Figure 2B). Our results suggest that similar to KCl stimulation, chronic SMS induced an immediate change in the structural plasticity of the AIS of cortical neurons that persisted after the stimulation period concluded.

## Discussion

Here, we demonstrate the novel finding that AIS structural plasticity of P1 derived inhibitory cortical neurons in cell culture can be induced with the chronic application of 15mM KCl or 0.5T SMS. KCl induced significant reductions in AIS length compared to control neurons immediately following 6 or 48 hours of stimulation, and these reductions persisted at 24 hours after return to control medium. These changes in AIS length were also accompanied by a significant distal relocation of the proximal start of the AIS after 48 hours of KCl treatment, which persisted for 24 hours following return to control medium. Similar to the changes induced by 15mM KCl, we found that 0.5T SMS delivered for 6 or 48 hours resulted in a significant reduction in AIS length. While this change persisted for 24 hours following cessation of 6 hours of SMS and return to control medium, the AIS returned to control lengths 24 hours following 48 hours of stimulation. A significant distal relocation of the AIS was observed after 48 hours of SMS which was present 24 hours after the stimulation had ended. Additionally, a delayed distal relocation was observed 24 hours following 6 hours of SMS.

### KCl induces AIS shortening and distal relocation in inhibitory neurons

Our results show that similar to studies using embryonically derived neurons in cell culture, sustained depolarisation of postnatally derived cortical neurons in cell culture by 15mM KCl induces a shortening and/or distal relocation of the AIS. Interestingly, we found that the AIS of inhibitory cortical neurons did not return to control lengths and locations after returning the neurons to control medium for 24 hours. This response has previously been observed in GABAergic dopaminergic interneurons taken from the olfactory bulb, where the AIS lengths did not return to control values in cell culture until 5 days post-stimulation (18). While our neurons are younger (relative to DIV) than those used in the previously mentioned study and may exhibit a slightly different developmental response in AIS structural plasticity (12, 15, 19), our results provide further evidence that structural AIS plasticity in inhibitory neurons is relatively slow to return to baseline. We speculate that this may be an intrinsic property of inhibitory neurons, irrespective of the brain regions where the cells are cultured from, but this requires further validation.

### SMS induces AIS plasticity reflective of increased neuronal activity

Similar to the KCl intervention, 0.5T SMS induced AIS structural plasticity in inhibitory neurons when delivered for 6 and 48 hours (i.e. chronic stimulation). Given that previous studies have shown that AIS shortening and distal relocation occurs following “excitatory” forms of stimulation (13, 18), this would suggest that chronic SMS is also a form of excitatory stimulation. This finding contrasts with the acute application of SMS (10-30min) which generally depresses cortical excitability and neural activity (2–7). Therefore, we speculate that the effect of SMS on neural activity switches from inhibition to excitation when using long durations. However, along with determining the functional consequence of the AIS structural plasticity induced on neuronal excitability (e.g., on action potential generation), further studies are needed to determine the mechanism of action of chronic SMS.

Unlike KCl stimulation, SMS displayed additional immediate and delayed effects on AIS structural plasticity which varied between 6 and 48 hours of SMS. In the 6-hour SMS protocol, there was an immediate and sustained shortening of the AIS length and delayed distal shift in AIS position. In contrast, the 48-hour SMS protocol results in an immediate and sustained distal shift in the AIS and only an immediate shortening of the AIS length. In short, this suggests that 6 hours of SMS is a more potent driver of AIS length changes whereas 48 hours of SMS is a more potent driver of AIS relocation. To our knowledge, the plasticity mechanisms that separate remodelling of the AIS length from location are not yet known. Therefore, we are unable to speculate why the two stimulation durations targeted different forms of AIS structural plasticity. Nevertheless, our results provide evidence that SMS duration is a key parameter in determining the type of structural AIS plasticity that is induced.

As an experimental tool *in vitro*, chronic SMS may provide a cheaper and simpler alternative to other methods that are used to drive AIS plasticity (e.g. optogenetics). However, *in vivo*, SMS is typically applied to humans within the range of 2 minutes −30 minutes in a single stimulation session (2, 5, 6, 9, 20). Whilst we have shown that 6 and 48 hours of stimulation induces AIS structural plasticity *in vitro*, the safety of such stimulation *in vivo* is unknown. Therefore, if chronic SMS was to be translated to drive AIS plasticity in the brain, its safety would have to be validated first.

In conclusion, we have demonstrated that chronic SMS can induce structural AIS plasticity in inhibitory cortical neurons derived from postnatal mice. The observed changes were similar to AIS plasticity induced with 15mM KCl stimulation with chronic SMS shortening the length of the AIS and location from the soma. Interestingly, some of the SMS-induced AIS structural plasticity was long lasting, persisting for 24 hours post stimulation. These findings provide a proof of principle that chronic SMS can induce neural plasticity and has the potential to be used as a non-invasive tool to drive AIS plasticity.

## Supplementary Figure

**Supplementary Figure 1.**
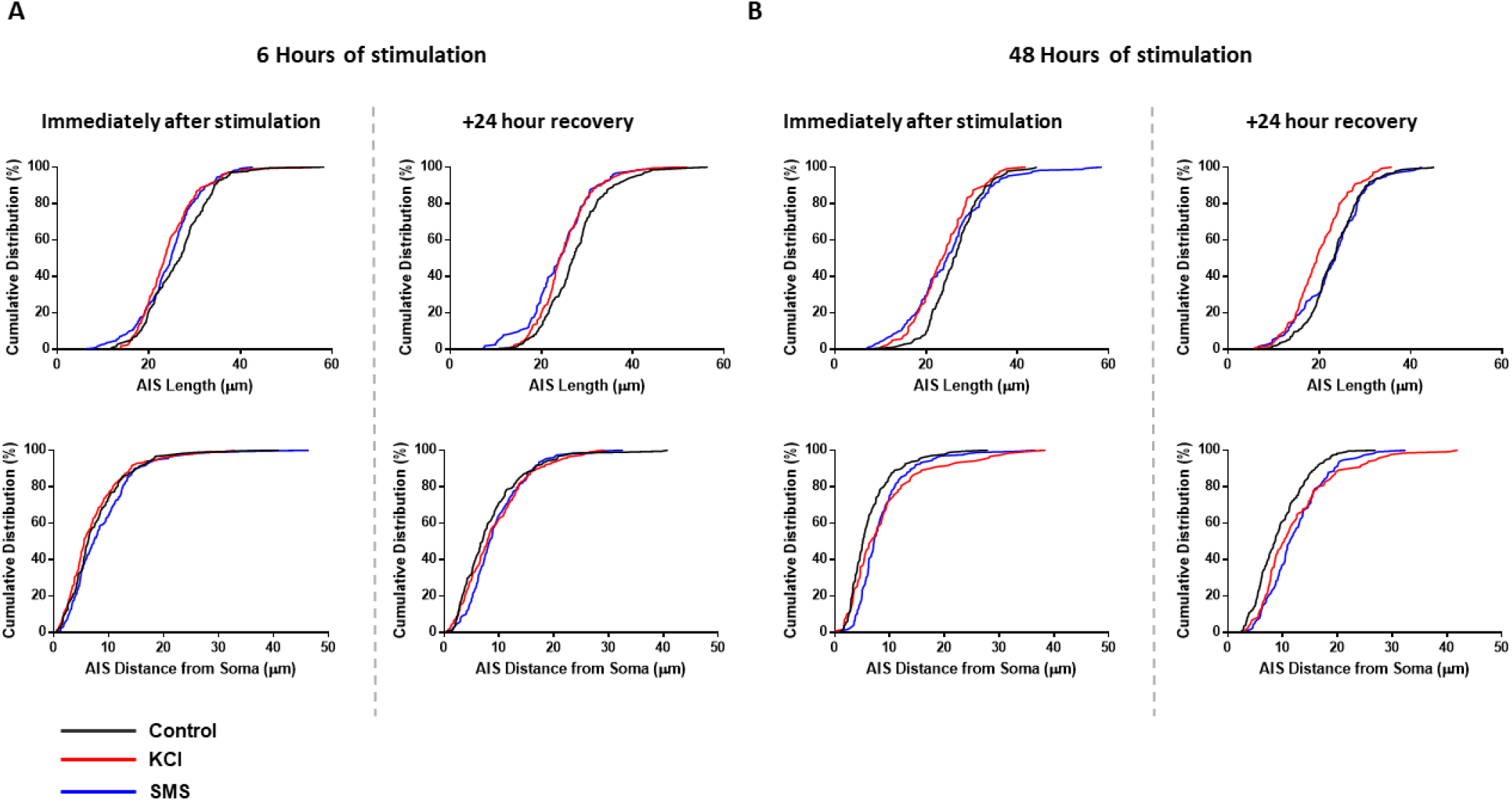
Cumulative distribution frequency plots of AIS length (top) and distance from soma (bottom) immediately after (left panel) and 24 hours after (right panel) 48 hours of stimulation.

## References

1. Nojima I, Oliviero A, Mima T. Transcranial static magnetic stimulation - From bench to bedside and beyond. Neuroscience Research. 2020;156:250–5.

2. Oliviero A, Mordillo-Mateos L, Arias P, Panyavin I, Foffani G, Aguilar J. Transcranial static magnetic field stimulation of the human motor cortex. The Journal of Physiology. 2011;589(20):4949–58.

3. Kirimoto H, Tamaki H, Matsumoto T, Sugawara K, Suzuki M, Oyama M, et al. Effect of transcranial static magnetic field stimulation over the sensorimotor cortex on somatosensory evoked potentials in humans. Brain Stimulation. 2014;7(6):836–40.

4. Dileone M, Mordillo-Mateos L, Oliviero A, Foffani G. Long-lasting effects of transcranial static magnetic field stimulation on motor cortex excitability. Brain Stimulation. 2018;11(4):676–88.

5. Silbert BI, Pevcic DD, Patterson HI, Windnagel KA, Thickbroom GW. Inverse correlation between resting motor threshold and corticomotor excitability after static magnetic stimulation of human motor cortex. Brain Stimulation. 2013;6(5):817–20.

6. Shibata S, Watanabe T, Yukawa Y, Minakuchi M, Shimomura R, Ichimura S, et al. Effects of transcranial static magnetic stimulation over the primary motor cortex on local and network spontaneous electroencephalogram oscillations. Scientific Reports. 2021;11(1):8261.

7. Aguila J, Cudeiro J, Rivadulla C. Effects of static magnetic fields on the visual cortex: reversible visual deficits and reduction of neuronal activity. Cerebral Cortex. 2014;26(2):628–38.

8. Davila-Pérez P, Pascual-Leone A, Cudeiro J. Effects of transcranial static magnetic stimulation on motor cortex evaluated by different TMS waveforms and current directions. Neuroscience. 2019;413:22–30.

9. Lozano-Soto E, Soto-León V, Sabbarese S, Ruiz-Alvarez L, Sanchez-del-Rio M, Aguilar J, et al. Transcranial static magnetic field stimulation (tSMS) of the visual cortex decreases experimental photophobia. Cephalalgia. 2017;38(8):1493–7.

10. Grubb MS, Shu Y, Kuba H, Rasband MN, Wimmer VC, Bender KJ. Short- and long-term plasticity at the axon initial segment. The Journal of Neuroscience. 2011;31(45):16049.

11. Yamada R, Kuba H. Structural and functional plasticity at the axon initial segment. Frontiers in Cellular Neuroscience. 2016;10(250).

12. Jamann N, Dannehl D, Lehmann N, Wagener R, Thielemann C, Schultz C, et al. Sensory input drives rapid homeostatic scaling of the axon initial segment in mouse barrel cortex. Nat Communications. 2021;12(1):23.

13. Grubb MS, Burrone J. Activity-dependent relocation of the axon initial segment fine-tunes neuronal excitability. Nature. 2010;465(7301):1070–4.

14. Kuba H, Oichi Y, Ohmori H. Presynaptic activity regulates Na+ channel distribution at the axon initial segment. Nature. 2010;465(7301):1075–8.

15. Gutzmann A, Erguel N, Grossmann R, Schultz C, Wahle P, Engelhardt M. A period of structural plasticity at the axon initial segment in developing visual cortex. Frontiers in Neuroanatomy. 2014;8.

16. Evans MD, Sammons RP, Lebron S, Dumitrescu AS, Watkins TBK, Uebele VN, et al. Calcineurin Signaling Mediates Activity-Dependent Relocation of the Axon Initial Segment. The Journal of Neuroscience. 2013;33(16):6950.

17. Höfflin F, Jack A, Riedel C, Mack-Bucher J, Roos J, Corcelli C, et al. Heterogeneity of the axon initial segment in interneurons and pyramidal cells of rodent visual cortex. Frontiers in Cellular Neuroscience. 2017;11(332).

18. Chand AN, Galliano E, Chesters RA, Grubb MS. A distinct subtype of dopaminergic interneuron displays inverted structural plasticity at the axon initial segment. The Journal of Neuroscience. 2015;35(4):1573.

19. Akter N, Fukaya R, Adachi R, Kawabe H, Kuba H. Structural and functional refinement of the axon initial segment in avian cochlear nucleus during development. The Journal of Neuroscience. 2020;40(35):6709.

20. Nojima I, Koganemaru S, Fukuyama H, Mima T. Static magnetic field can transiently alter the human intracortical inhibitory system. Clinical Neurophysiology. 2015;126(12):2314–9.

